# *rearrvisr*: an R package to detect, classify, and visualize genome rearrangements

**DOI:** 10.1101/2020.06.25.170522

**Authors:** Dorothea Lindtke, Sam Yeaman

## Abstract

The identification of genome rearrangements is of direct relevance for understanding their potential impacts on evolution and disease. However, available methods that detect or visualize rearrangements from deviations in gene order do not map them onto a genome of interest, complicating downstream analysis. In this work, we present *rearrvisr*, an R package that implements a novel algorithm for the identification and classification of rearrangements. In contrast to other software, it projects rearrangements onto a single genome, facilitating the localization of rearranged regions and estimation of their extent. We show that our tool achieves high precision and recall scores on simulated data, and illustrate the utility of our method by applying it to a data set generated from publicly available *Drosophila* genomes. The package is freely available from GitHub (https://github.com/dorolin/rearrvisr) and can be installed directly from R.

## 1 Introduction

Chromosomal rearrangements are a significant source of genetic variation (Bhutkar *et al.*, 2008; The 1000 Genomes Project Consortium, 2015) and play an important role in evolution and disease (Lisch, 2013; Rieseberg, 2001; Weischenfeldt *et al.*, 2013). Rearrangements are commonly detected through the pairwise alignment of genomes or long-read sequences (Frith and Kahn, 2018; Soderlund *et al.*, 2011), or inferred from anomalies when short-read sequences are mapped to a reference genome (Tattini *et al.*, 2015). However, read mapping anomalies provide only suboptimal, indirect evidence for rearrangements and are difficult to visualize, while current software for the identification and visualization of rearrangements from pairwise alignments (e.g., Mauve; Darling *et al.*, 2004; SyMAP; Soderlund *et al.*, 2011) or through computation of an optimal scenario of rearrangement events (e.g., GRIMM; Tesler, 2002; UniMoG; Hilker *et al.*, 2012) do not allow an easy localization of rearrangements along a genome of interest (i.e., a “focal” genome). Further, the detection of structural genomic changes that occurred during the evolution of a particular lineage requires the reconstruction of an ancestral genome, but available methods do not project rearrangements onto a focal genome (e.g., ANGES; Jones *et al.*, 2012; MGRA; Avdeyev *et al.*, 2016; MLGO; Hu *et al.*, 2014).

To overcome the shortcomings of available software, we developed the R package *rearrvisr* to identify, classify, and visualize rearrangements for a focal genome relative to an ancestral genome reconstruction or an extant genome (i.e., a “compared” genome). In contrast to other software, our method directly maps rearrangements onto the focal genome, enabling the localization of rearranged genomic regions and facilitating the determination of their extent. This functionality is of significance, for example, for correlational studies or for identifying highly rearranged genomic regions. The use of the platform-independent programming environment R (R Core Team, 2018) allows an easy integration in custom analysis. The package can take as input chromosome-level or fragmented genome representations in form of maps that specify the order and orientation of markers. The algorithm then searches for deviations in marker order and orientation in the focal genome relative to the compared genome to detect and classify rearrangements. In the following, we describe the basics of our algorithm, outline the functionalities of the package, and apply it to simulated and real data. More details are available in the Supplementary information. Function documentation, a tutorial, and example files are included in the package.

## 2 Methods

### 2.1 Algorithm design

Consider two genomes, a *focal genome* (i.e., a genome for which rearrangements will be identified) and a *compared genome* (i.e., a genome that serves as a reference). Each genome consists of a set of *chromosomes* that contain contiguous sets of unique *markers* (i.e., genes, synteny blocks, or locally conserved segments of sequence). Common classes of rearrangements are fusions, fissions, reciprocal translocations, transpositions, and inversions, resulting from the movement or reversal of markers between or within chromosomes. They are detectable as differences in the order or orientation of orthologous markers in the sample genome (i.e., the focal genome) relative to the reference genome (i.e., the compared genome).

Currently available genome assemblies rarely represent complete sets of contiguous markers over whole biological chromosomes but instead are a collection of *genome segments* (*GSs*; i.e., scaffolds, super-scaffolds, or pseudo-chromosomes) that link smaller *contigs* (i.e., contiguous sets of ordered and oriented markers) together (Fig. 1A). We define *focal genome segments* (*FGSs*) as GSs that consist of an ordered and oriented set of contigs, where gaps between contigs arising from unlocalized or unplaced contigs during assembly are permitted (such gaps are commonly represented by stretches of N’s on an assembled scaffold). These unlocalized or unplaced contigs, which are biologically part of a larger GS, may be represented as an independent FGS in the focal genome. We define *compared genome segments* (*CGSs*) as contiguous sets of contigs without gaps, where markers that are biologically part of a GS are allowed to be missing but, in contrast to FGSs, cannot be located on an independent CGS. These CGSs define the order and orientation of markers prior to any rearrangement. CGSs correspond to the definition of ‘Contiguous Ancestral Regions’ (CARs) in Ma *et al.* (2006) and Chauve and Tannier (2008).

**Figure 1:**
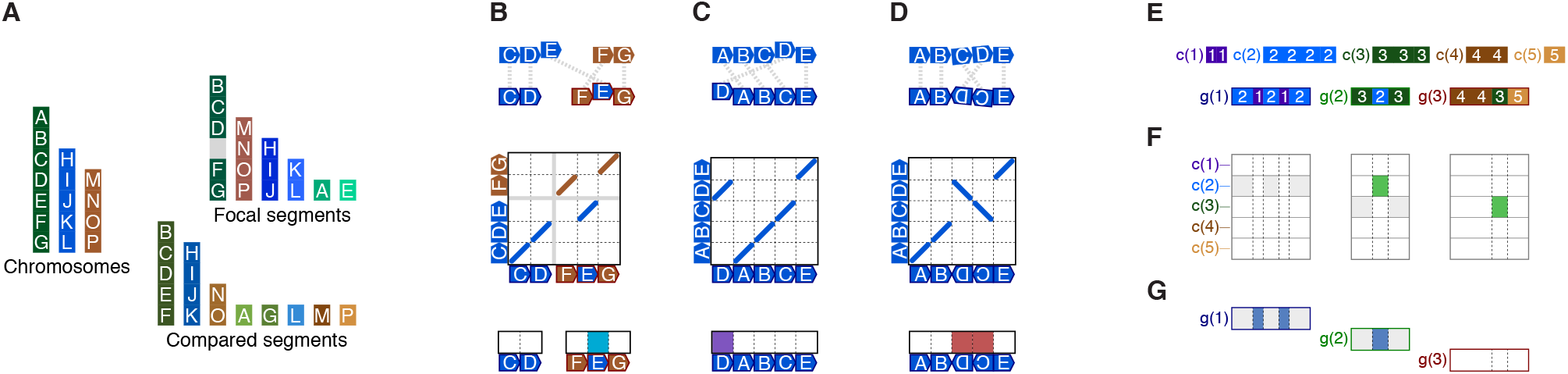
Genome segments and classes of rearrangements. **(A)** Three whole biological chromosomes are shown as colored bars on the left. Markers on each chromosome are uniquely labeled with letters. Focal genome segments (FGSs) representing the chromosomes on the left are shown on the top right. In this example, not all markers are placed on three super-scaffolds and instead remain on separate minor scaffolds. The gray rectangle represents a gap between markers D and F, and marker E that corresponds to this gap is located on a separate minor scaffold. Compared genome segments (CGSs) representing the chromosomes on the left are shown on the bottom right. In this example, the differences between FGSs and CGSs only arise from their assembly, not from rearrangements. **(B-D)** Classes of rearrangements. The top row illustrates dual synteny plots between CGSs (top) and FGSs (bottom). Oriented markers are shown as pentagons, and orthologs are connected with dotted lines. The middle row gives pairwise dotplots. The bottom row illustrates how rearrangements are mapped on FGSs in the *rearrvisr* output. **(B)** Nonsyntenic moves. **(C)** Syntenic moves. **(D)** Inversions. **(E-G)** Detection of nonsyntenic moves in *rearrvisr*. **(E)** IDs of five CGSs, c(i), are mapped on three FGSs, *g*(*j*). CGS IDs are indicated with integers. **(F)** Detection of nonsyntenic moves class I across FGSs, separately for each CGS, illustrated with different rows per c(i). Columns correspond to orthologs in *g*(*j*). Filled cells indicate markers where the contiguity of a *c*(*i*) is disrupted by mapping blocks of *c*(*i*) onto different FGSs *g*(*j*) and *g*(*d* ≠ *j*). Colored cells indicate blocks that are more parsimonious to have moved. **(G)** Detection of nonsyntenic moves class II across CGSs, separately for each FGS. Filled cells indicate markers where the contiguity of one or more *c*(*i*) is disrupted within a *g*(*j*) by insertions of blocks from other CGSs *c*(*u* ≠ *i*). Colored cells indicate blocks that are more parsimonious to have moved.

We define a *genome G* as a set of *N*_*G*_ GSs, *G* = {*g*(1), …, *g*(*N*_*G*_)}. The order of GSs within a genome is irrelevant. Each GS *g*(*p*) contains *n*_*p*_ markers, 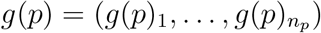, 1 ≤ *p* ≤ *N*_*G*_. The ends of a GS (e.g., the telomeres) are denoted as *g*(*p*)_0_ and 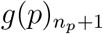, and requiring 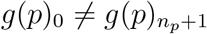 (i.e., circular GSs are not permitted). Each marker has two *adjacencies*, one at its *head* and one at its *tail* (e.g., the head of marker *g*(*p*)_*i*+1_ is adjacent to the tail of marker *g*(*p*)_*i*_, 1 ≤ *i* < *n*_*p*_, the head of marker *g*(*p*)_1_ is adjacent to *g*(*p*)_0_, and the tail of marker 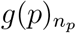 is adjacent to 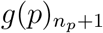). Each marker has its *marker orientation* (i.e., strand) given by a + or − sign (the + is omitted for better readability), where marker −*g*(*p*)_*i*_ has its head and tail reversed relative to marker *g*(*p*)_*i*_. A GS can be read in *ascending* (i.e., standard) or in *descending* (i.e., reversed) direction (e.g., 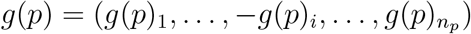 is equivalent to 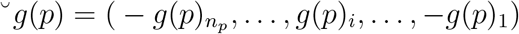). As a simplification, the descriptions below assume that CGSs are ascending and only contain positively oriented markers.

We define a *rearrangement* as the formation of a novel adjacency of marker *c*(*q*)_*m*_ from CGS *c*(*q*) that results in the disruption of the contiguity of markers on CGS *c*(*q*) (Supplementary Fig. S1). As a clear distinction between inter- and intrachromosomal rearrangements is not possible for fragmented genome assemblies, we here classify rearrangements as nonsyntenic or syntenic moves, or as inversions (Fig. 1B-E; defined below). Nonsyntenic and syntenic moves approximate inter- and intrachromosomal transpositions, respectively, on biological chromosomes. Fusions and fissions will not result in patterns inconsistent with fragmented genome assemblies and are therefore not considered.

Let an *internal boundary* of CGS *c*(*q*) on GS *g*(*p*) (i.e., a FGS or its ancestor) be an adjacency between two markers on *g*(*p*), *g*(*p*)_*i*_ and *g*(*p*)_*i*+1_, 1 ≤ *i* < *n*_*p*_, where marker *g*(*p*)_*i*_ has an ortholog in CGS *c*(*q*) and marker *g*(*p*)_*i*+1_ and ortholog in CGS *c*(*u*), *u* ≠ *q*. Further, an *end boundary* of *c*(*q*) on *g*(*p*) exists if *g*(*p*)_1_ or 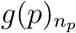 has an ortholog in *c*(*q*). We define a *nonsyntenic move* as the disruption of the contiguity of *c*(*q*) by the formation of a novel adjacency between the tail of a marker *c*(*q*)_*m*_, *m* < *n*_*q*_, having an ortholog *g*(*p*)_*i*_ on *g*(*p*), and a marker *g*(*p*)_*i*+1_ on *g*(*p*), leading to the formation of a novel internal boundaries of *c*(*q*) on *g*(*p*) or a different GS *g*(*d*), *d* ≠ *p* (Supplementary Fig. S1A-C). A nonsyntenic move may result from the movement of a block of orthologs [*c*(*q*)_*l*_, …, *c*(*q*)_*m*_], 1 ≤ *l* ≤ *m* < *n*_*q*_, from *g*(*p*) onto *g*(*d*) (Supplementary Fig. S1B). It may also result from the insertion of a block of orthologs [*c*(*u*)_*s*_, …, *c*(*u*)_*t*_], 1 ≤ *s* ≤ *t* ≤ *n*_*u*_, from *g*(*d*) between orthologs *c*(*q*)_*m*_ and *c*(*q*)_*m*+1_, 1 ≤ *m* < *n*_*q*_, on *g*(*p*) (Supplementary Fig. S1C). Analogous definitions apply for the formation of a novel adjacency of the head of marker *c*(*q*)_*l*_, 1 < *l* ≤ *m* ≤ *n*_*q*_. We define a *syntenic move* as the disruption of the contiguity of *c*(*q*) by the formation of a novel adjacency between a marker *c*(*q*)_*k*_ and a marker *c*(*q*)_*m*_ on the same *g*(*p*) by swapping the positions of two blocks [*c*(*q*)_*k*_, …, *c*(*q*)_*l*−1_] and [*c*(*q*)_*l*_, …, *c*(*q*)_*m*_], 1 ≤ *k* < *l* ≤ *m* ≤ *n*_*q*_, without formation of additional boundaries (Supplementary Fig. S1D). We define an *inversion* as the disruption of the contiguity of *c*(*q*) by the formation of a novel adjacency between marker *c*(*q*)_*l*−1_ and the tail of marker *c*(*q*)_*m*_, 1 < *l* ≤ *m* ≤ *n*_*q*_, by reversing a block [*c*(*q*)_*l*_, …, *c*(*q*)_*m*_] to [−*c*(*q*)_*m*_, …, −*c*(*q*)_*l*_] on the same *g*(*p*), without formation of additional boundaries (Supplementary Fig. S1E). The definition is analogous for a novel adjacency of the head of marker *c*(*q*)_*l*_, 1 ≤ *l* ≤ *m* < *n*_*q*_.

#### 2.1.1 General principle

Let the order of markers in the compared genome be uniquely encoded by a hierarchical set of indices, one set of two positive integers per marker, referred to as *CGS ID* and *element ID*. The CGS ID is unique to each CGS, and the element ID is strictly increasing with the reading direction of markers on a CGS. Markers without orthologs are excluded, and remaining markers in the compared genome are associated (i.e., mapped) to their orthologs in the focal genome, potentially altering the original sequence of their indices when read by their order in the focal genome (Fig. 1E). Rearrangements are expected to lead to inconsistencies in the altered sequence of indices, corresponding to disruptions of the contiguity of a CGS. These inconsistencies are detected in a stepwise process. First, nonsyntenic moves are identified through the detection of an unsupported number of shifts between CGS IDs (i.e., boundaries) along the focal genome, which signify inconsistencies in CGS IDs. Then, syntenic moves and inversions are identified through a discrepancy in the expected order of element IDs, where a discrepancy in the expected order is given when mapped element IDs are neither strictly increasing nor decreasing. In both steps, markers that are more parsimonious to have caused an observed inconsistency are identified as rearranged. The two steps are briefly described in the following. More details and pseudocode are available in the Supplementary information.

#### 2.1.2 Nonsyntenic moves

For fragmented genome assemblies, multiple CGSs may be mapped onto one FGS, or a CGS may be split across multiple FGSs (Fig. 1E). Thus, the presence of shifts in CGS IDs along an FGS indicates a disruption of the contiguity of a CGS only when they result in an excess number of internal boundaries. Briefly, while for a CGS any number of end boundaries across one or more FGS (without internal boundaries) or two internal boundaries on one FGS (without end boundaries) are consistent with a fragmented genome assembly without nonsyntenic moves, three or more internal boundaries or certain arrangements of two internal boundaries with end boundaries are not. For example, a nonsyntenic move of a block of markers with orthologs on CGS *c*(*q*) from GS *g*(*p*) onto GS *g*(*d*) will result in one or two additional internal boundaries on *g*(*d*), by maintaining the already existing internal boundaries (if any). We detect this type of rearrangement, which we term *class I nonsyntenic move*, by considering the number and arrangement of internal and end boundaries across all *g*(*j*), 1 ≤ *j* ≤ *N*_*G*_, separately for each *c*(*i*), 1 ≤ *i* ≤ *N*_*C*_ (Fig. 1F). Similarly, a nonsyntenic move of a block of markers with orthologs on CGS *c*(*u*) from GS *g*(*d*) onto GS *g*(*p*) will result in two additional internal boundaries on genome segment *g*(*p*). We detect this type of rearrangement, which we term *class II nonsyntenic move*, by considering the number and arrangement of internal and end boundaries across all *c*(*i*), 1 ≤ *i* ≤ *N*_*C*_, separately for each *g*(*j*), 1 ≤ *j* ≤ *N*_*G*_ (Fig. 1G). Although in many cases nonsyntenic moves will belong to both classes, the difference between them is that class I corresponds to a disruption of the contiguity of *c*(*i*) by splitting its markers *between* two (or more) *g*(*p*) and *g*(*d*), *p* ≠ *d*, while class II corresponds to a disruption of the contiguity of *c*(*i*) by the insertion of markers from *c*(*u*), *u* ≠ *i*, *within* a single *g*(*j*). If *c*(*i*) and *c*(*u*) are located on the same biological chromosome, class II is equivalent to a syntenic move. If *g*(*p*) and *g*(*d*) are located on the same biological chromosome, class I is equivalent to a fusion of *c*(*i*) and *c*(*u*), followed by a syntenic move. The detection of class I nonsyntenic moves takes into account that FGSs are not permitted to form circles to link their ends along matching CGSs, and that small FGSs may represent unplaced contigs in the focal genome assembly, rather than rearrangements. Nonsyntenic moves that are indistinguishable from fragmented genome assemblies based on their number and arrangement of boundaries are ignored.

The positions of markers involved in nonsyntenic moves are determined and assigned with *tags* (i.e., values between 0 and 1). Any nonsyntenic move involves two blocks of CGS *c*(*q*), [*c*(*q*)_*l*_, …, *c*(*q*)_*m*_] and 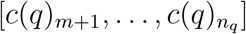, 1 ≤ *l* ≤ *m* < *n*_*q*_, and potentially a third block from GS *g*(*p*), [*g*(*p*)_*i*_, …, *g*(*p*)_*j*_], 1 ≤ *i* ≤ *j* ≤ *n*_*p*_, that inserted between these blocks. As the true rearrangement scenario is unknown and blocks are moved only relative to each other in the observational data, the principle of parsimony is applied to specify comparatively few markers as being rearranged (i.e., based on the relative number of markers per block). The problem of defining the positions of moved markers is further addressed by assigning different tag values to markers that are part of rearranged blocks, where larger values indicate blocks that are more parsimonious to have been moved, relative to alternative blocks. Full details are available in the Supplementary information.

#### 2.1.3 Syntenic moves and inversions

Rearrangements between two markers located on the same CGS *c*(*q*) and with orthologs on the same FGS *g*(*p*) that do not result in the formation of novel boundaries (i.e., syntenic moves and inversions) can be identified through a discrepancy in the expected order of element IDs. Let *c*(*q*)_*m*_ and *c*(*q*)_*n*_ be markers on CGS *c*(*q*), 1 ≤ *m* < *n* ≤ *n*_*q*_, with orthologs *g*(*p*)_*i*_ and *g*(*p*)_*j*_, respectively, on FGS *g*(*p*), and 1 ≤ *i* < *j* ≤ *n*_*p*_. Further, let *ϕ*(*g*(*p*)_*i*_) and *ϕ*(*g*(*p*)_*j*_) be the element IDs of markers *c*(*q*)_*m*_ and *c*(*q*)_*n*_, respectively, read by their order on *g*(*p*). A disruption of the contiguity of *c*(*q*), aligned in ascending direction to *g*(*p*), is then identified if *ϕ*(*g*(*p*)_*i*_) > *ϕ*(*g*(*p*)_*j*_) (as the expected order is *ϕ*(*g*(*p*)_*i*_) < *ϕ*(*g*(*p*)_*j*_) for an ascending alignment of *c*(*q*)). Similarly, a disruption of the contiguity of −*c*(*q*) (i.e., aligned in descending direction to *g*(*p*)) is identified if *ϕ*(*g*(*p*)_*i*_) < *ϕ*(*g*(*p*)_*j*_) (as then the expected order is *ϕ*(*g*(*p*)_*i*_) > *ϕ*(*g*(*p*)_*j*_)). As a simplification, the descriptions below are restricted to CGSs that are aligned in ascending (i.e., standard) direction to *g*(*p*).

Either a syntenic move (i.e., the interchange between two blocks, one containing *c*(*q*)_*m*_, and one containing *c*(*q*)_*n*_) or an inversion (i.e., the reversal of one block containing both *c*(*q*)_*m*_ and *c*(*q*)_*n*_) will result in *ϕ*(*g*(*p*)_*i*_) > *ϕ*(*g*(*p*)_*j*_). As two GSs containing the same set of orthologs can be transformed into each other either by block interchanges, reversals, or a combination of both (for a review see Li *et al.*, 2006), rearrangements are classified as syntenic moves or inversions based on the principle of parsimony to reduce the number of required rearrangements and involved markers. The main steps applied for identifying syntenic moves and inversions are briefly outlined below. More details are available in the Supplementary information.

First, the set of markers on *g*(*p*) that have orthologs on *c*(*q*), *σ*_*pq*_, are identified and their element IDs *ϕ*(*g*(*p*)_*i*_), *g*(*p*)_*i*_ ∈ *σ*_*pq*_, are iteratively clustered into blocks of local synteny. For this, *ϕ*(*σ*_*pq*_) are ranked and clustered into a set of blocks *B*(*λ*). Each block *B*(*λ*)_*b*_ ∈ *B*(*λ*) groups a sequence of ranks that either differ exactly by 1 or by −1. Blocks are then assigned new ranks according to the values of their contained ranks, and the clustering of blocks into larger blocks based on their ranks is iterated over clustering steps *λ*. Clustering terminates at the ultimate step 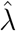 where no additional clustering can be achieved. At each step 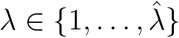, and for each block *B*(*λ*)_*b*_ at step *λ*, the *block orientation o*(*B*(*λ*)_*b*_) is determined, which can be ascending (ranks are increasing), descending (ranks are decreasing), or missing if a block is formed by a single rank. At any *λ*, a partitioning of ranks into more than one block points to a discrepancy in the expected order of element IDs and thus a rearrangement.

Second, if clustering of ranks into a single ultimate block was not possible, *complex* rearrangements are identified. This can be the case, for example, when multiple syntenic moves have crossed over in *σ*_*pq*_. A small set of blocks 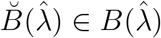 is then determined, based on the principle of parsimony, that needs to be removed to achieve final clustering of remaining blocks into a single block after further clustering iterations. Each block in 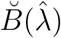 is classified as a syntenic move and markers part of it are tagged with a value of 1. The ultimate clustering step after removal of blocks in 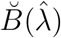 defines the new 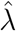.

Then, rearrangements are identified and classified as syntenic moves or inversions based on the orientation and number of blocks at clustering steps *λ*. Let the *higher-level block* 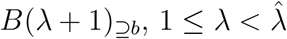, be the block at step *λ* + 1 that contains block *B*(*λ*)_*b*_, and let the *lower-level blocks s*(*b*) = (*s*(*b*)_1_, …, *s*(*b*)_*n_b_*_) be the blocks at step *λ* − 1 that are contained in 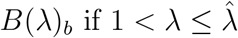, or the ranks of element IDs that were initially clustered into *B*(*λ*)_*b*_ if *λ* = 1. Further, let the marker *directionality* of *g*(*p*)_*i*_ be defined as identical or inverted if orthologs *g*(*p*)_*i*_ and *c*(*q*)_*m*_ ∈ *σ*_*pq*_ have the same or opposing marker orientation, respectively.

To identify and classify rearrangements, the *alignment direction O*(*σ*_*pq*_) for the joint set of markers *c*(*q*)_*m*_ ∈ *σ*_*pq*_ to *g*(*p*) is determined, which can be ascending (i.e., standard alignment) or descending (i.e., inverted alignment). *O*(*σ_pq_*) is defined as the orientation of the final block 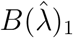, 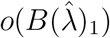. if 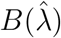 contains three or more lower-level blocks. Otherwise, if 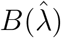 contains exactly two lower-level blocks, and both have reversed orientation (or inverted marker directionality if 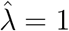 to 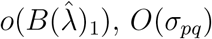 is set to the reverse of 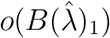, and a syntenic move between the two lower-level blocks is identified. This measure reduces the number of required rearrangements, as without reversal of *O*(*σ*_*pq*_) two rearrangements would be assigned below.

Subsequently, rearrangements are identified and classified for each clustering step 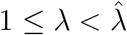. Let the *higher-level orientation* of block *B*(*λ*)_*b*_, 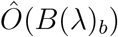, be defined as *O*(*σ*_*pq*_) for 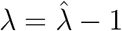, and by the block orientation and rearrangement type (if any; below) of *B*(*λ* + 1)_⊇*b*_ for 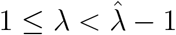. Starting with blocks from the penultimate iteration, 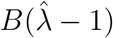, blocks with orientation *o*(*B*(*λ*)_*b*_) reversed relative to 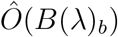 are identified as syntenic moves or inversions. All blocks *B*(*λ*)_*b*_ at each clustering step are tested for rearrangements by stepwise decrementing *λ* by 1, from 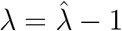 to *λ* = 1. The rearrangement class is parsimoniously assigned based on the number of lower-level blocks and their orientation (or marker directionalities for *λ* = 1). If a syntenic move was assigned to *B*(*λ*)_*b*_, 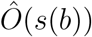 is set to the reverse of *o*(*B*(*λ*)_*b*_).

As a final step, markers *g*(*p*)_*i*_ ∈ *σ*_*pq*_ are tested for single-marker inversions. A *single-marker inversion* is identified if the directionality of *g*(*p*)_*i*_ deviates from its expected directionality. The expected directionality of *g*(*p*)_*i*_ is computed based on *O*(*σ*_*pq*_) and the number of inversion events that were assigned to *g*(*p*)_*i*_ across steps 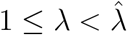. Markers that are part of single- or multi-marker inversions are tagged with a value of 1. For syntenic moves, larger tag values are assigned to markers that are part of the block that is more parsimonious to have been moved, relative to the alternative block.

#### 2.1.4 Syntenic moves and inversions with marker order uncertainty

CGSs can optionally be represented by *PQ-trees*, combinatorial structures consisting of *leaves*, *P-nodes*, and *Q-nodes*, which can incorporate ambiguity in marker order (Booth and Lueker, 1976; Chauve and Tannier, 2008). Briefly, each marker is represented by a *leaf*, which is connected through a hierarchical sequence of *nodes* (*P-nodes* or *Q-nodes*) to the *root* of the *PQ-tree* (i.e., the top-level node, representing a CGS). The type of the node specifies the order of its children: they are in arbitrary order for *P-nodes*, and in fixed order (including their reversal) for *Q-nodes*. *Branches* connect a node (including the root) to its children (i.e., lower-level nodes or leaves), and are labeled according to the position of the children on a node with unique and strictly increasing positive integers referred to as *branch positions*. The position of each marker *c*(*q*)_*m*_ on CGS *c*(*q*) can thus be precisely described by the sequence of branch positions traversed across all hierarchy levels of a *PQ-tree*. Without marker order ambiguity, *c*(*q*) can be represented by a single *Q-node* at the root with branches directly connecting to the leaves, where branch positions correspond to element IDs.

As before with element IDs, the set of branch positions for marker *c*(*q*)_*m*_ is associated (i.e., mapped) to its ortholog in the focal genome, *g*(*p*)_*i*_. The algorithm then detects syntenic moves and inversions hierarchically, from the root to the leaves of the *PQ-tree*, by generating subsets of markers in *g*(*p*) for each node on each hierarchy level. Each subset is then tested separately for rearrangements by searching for inconsistencies in the sequence of their branch positions at the respective hierarchy level, thereby allowing for uncertainty in the order of markers in *c*(*q*). For *Q-nodes*, the assignment of syntenic moves and inversions follows steps described above for element IDs, with slight adjustments accounting for the fact that branch positions, unlike element IDs, are not necessarily unique for a given hierarchy level. For *P-nodes*, all rearrangements are assigned as syntenic moves and are detected by steps that follow the principle of identifying class II nonsyntenic moves. More details are available in the Supplementary information.

### 2.2 Implementation and usage

Our algorithm is implemented in the R package *rearrvisr*. Two files that specify the location and orientation of markers along each genome are required as input. The compared genome can optionally incorporate ambiguity in marker order (e.g., it can be provided as a genome reconstruction from the software ANGES; Jones *et al.*, 2012). Markers can be any orthologous genomic regions that are unique within each genome, such as genes, synteny blocks, or locally conserved segments of sequence (e.g., identified with software such as OMA, Altenhoff *et al.*, 2015, or DAGchainer, Haas *et al.*, 2004). The package includes example data as well as functions to convert and verify input files (the `convertPQtree()`, `genome2PQtree()`, and `checkInfile()` functions).

Nonsyntenic moves, syntenic moves, and inversions (Fig. 1) are identified with the `computeRearrs()` function. The output includes tables indicating rearranged markers, the positions of breakpoints, and additional information (e.g., on the alignment direction). The `summarizeBlocks()` function summarizes these tables for each conserved block of markers. The `genomeImagePlot()` function visualizes rearranged markers along their genomic position with different colors per rearrangement class (Fig. 2B), allowing easy visual assessment of location and extent of rearranged regions. The `genomeRearrPlot()` function shows the alignment of conserved blocks to the focal genome and visualizes all (potentially nested) rearrangements (Fig. 2C), facilitating the interpretation of the results and enabling the derivation of alternative rearrangement assignments. A tutorial is available in the package vignette.

**Figure 2:**
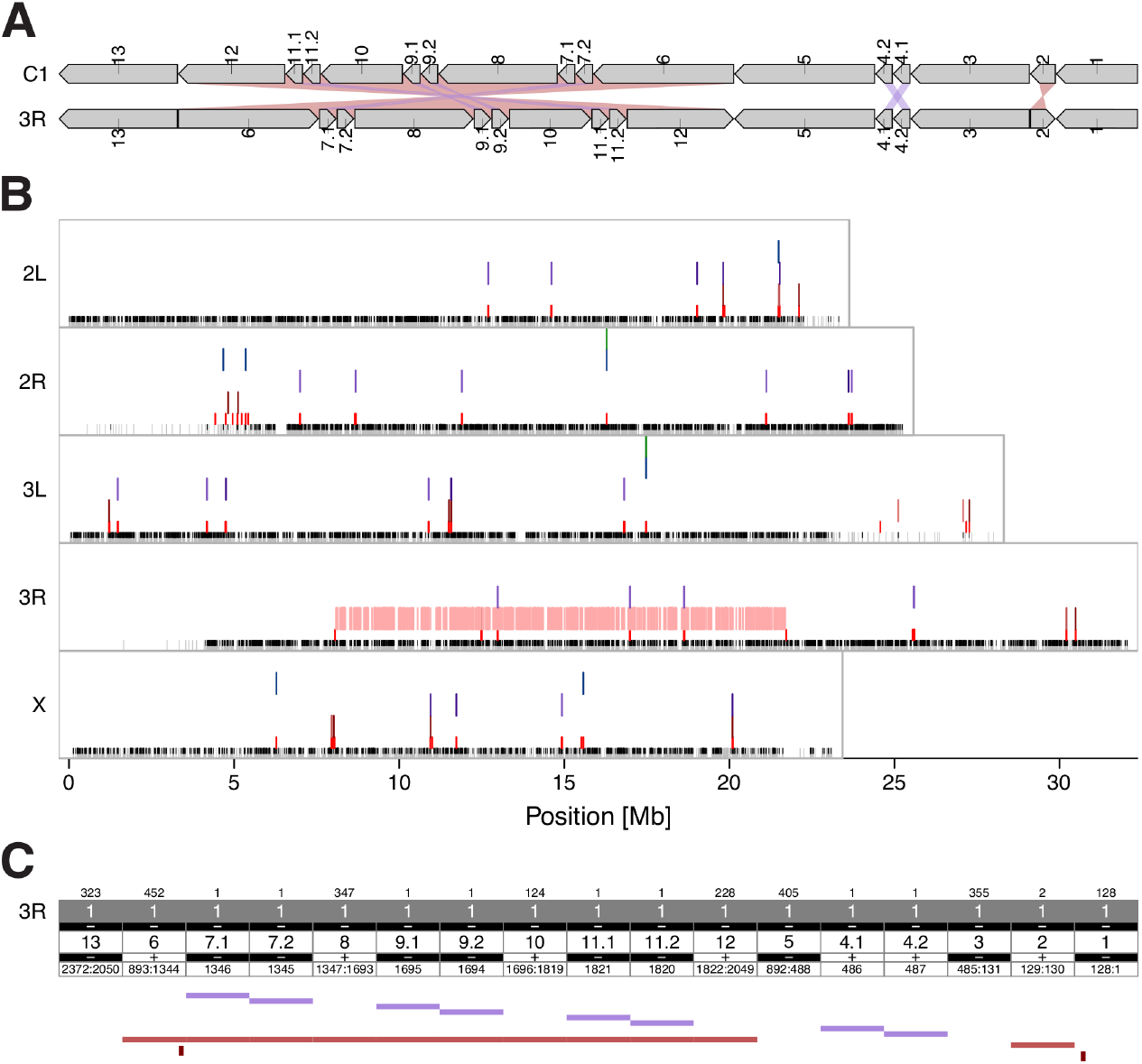
Rearrangements in *D. melanogaster* that occurred after divergence from the *melanogaster* subgroup ancestor were identified and visualized with *rearrvisr*. **(A)** Schematic of the alignment between reconstructed ancestral genome segment C1 and *D. melanogaster* chromosome 3R. Pentagons illustrate conserved marker blocks and their orientation. Purple and maroon bands highlight syntenic moves and inversions. **(B)** Rearranged markers are indicated by colored vertical lines at their genomic position for five *D. melanogaster* chromosomes on separate rows per rearrangement class. The two bottom rows show rearrangement breakpoints in red and positions of orthologous markers in black. **(C)** Information for each conserved block is in columns, with the number of contained markers indicated on top. The upper part illustrates the alignment of ancestral genome segment C1 to *D. melanogaster* chromosome 3R, and colored bars at the bottom indicate rearranged blocks.

### 2.3 Performance measurements

To assess the performance of *rearrvisr* and compare it to other software, we simulated inter- and intrachromosomal transpositions (i.e., corresponding to nonsyntenic and syntenic moves, respectively), and inversions. The compared genome consisted of five chromosomes of equal size and a total of 5000 markers. The focal genome was generated by a fixed number of rearrangement events that occurred in random order and position on the compared genome. We simulated different sizes of rearranged blocks and different numbers of rearrangement events comprising an equal number of nonsyntenic moves, syntenic moves, and inversions. For some simulations, additional macro-rearrangements (i.e., one reciprocal translocation, or one fission and one fusion) were added at random time points to the simulations. We subsequently fragmented one or both genomes for a subset of simulations by adding 20 chromosome splits. For each setting 100 replicates were run. Further details are in the Supplementary information.

We compared *rearrvisr* to GRIMM 2.01 (Tesler, 2002) and UniMoG (Hilker *et al.*, 2012) on simulated data and *Drosophila* genomes. GRIMM and UniMoG are designed to compute an optimal scenario of rearrangement events required to transform one genome into another, using lists that specify location and orientation of markers along chromosomes as input. GRIMM, which is based on the Hannenhalli and Pevzner model (Hannenhalli and Pevzner, 1995), determines the minimal possible number of rearrangement steps by allowing inversions, reciprocal translocations, fusions, and fissions as rearrangement operations. It further assigns a corresponding sequence of operations, and computes the number of internal and external breakpoints. UniMoG, which implements the double cut and join (DCJ) model (Bergeron *et al.*, 2006; Yancopoulos *et al.*, 2005), additionally allows circularizations and decircularizations as rearrangement operations. As both programs do not identify rearranged markers, we instead compared the number of detected rearrangements between GRIMM, UniMoG, and *rearrvisr*, and additionally the relative frequency of different rearrangement classes and number of breakpoints between GRIMM and *rearrvisr* (these metrics are not output by UniMoG). For simulated data, the true number of breakpoints was approximated by the number of expected breakpoints per simulated rearrangement class, providing an upper bound as some breakpoints may fall on identical positions. We ran GRIMM with the default model and UniMoG with standard DCJ sorting. We ran *rearrvisr* under default settings and computed the number of identified rearrangements and breakpoints with the `summarizeRearrs()` function.

### 2.4 Application to *Drosophila*

We downloaded publicly available peptide sequences and annotation information for genes from 12 *Drosophila* species from Ensembl Release 91 (http://dec2017.archive.ensembl.org, *D. melanogaster*) or Ensembl Metazoa Release 37 (http://oct2017-metazoa.ensembl.org; *D. ananassae*, *D. erecta*, *D. grimshawi*, *D. mojavensis*, *D. persimilis*, *D. pseudoobscura*, *D. sechellia*, *D. simulans*, *D. virilis*, *D. willistoni*, and *D. yakuba*). We identified orthologs with OMA standalone v2.2.0 (Altenhoff *et al.*, 2015), and reconstructed the ancestral genome of the *melanogaster* subgroup (i.e., separating *D. melanogaster*, *D. simulans*, *D. sechellia*, *D. yakuba*, and *D. erecta* from the remainder of the *Drosophila* species), with ANGES v1.01 (Jones *et al.*, 2012). The ancestral genome comprised 8,973 orthologs, which were organized into 20 CARs (i.e., CGSs), encoded as *PQ-trees*. Further details on data preparation are in the Supplementary information.

We identified rearrangements and breakpoints with *rearrvisr*, GRIMM, and UniMoG by using the genome of *D. melanogaster* as focal and the genomes of *D. simulans* or *D. yakuba* as compared genome. In addition, we ran *rearrvisr* by using the genome of *D. melanogaster* as focal and the ancestral genome reconstruction as compared genome. To facilitate comparison, only orthologs located on the five main chromosome arms (2L, 2R, 3L, 3R, and X) and common to all four tested genomes were retained (8382 orthologs, on seven CARs).

## 3 Results

### 3.1 Performance evaluation

We assessed the performance of *rearrvisr* in detecting rearranged markers using simulated data by varying the total number of rearrangement events and sizes of rearranged blocks. Performance was evaluated separately for each rearrangement class by recall (calculated as the ratio of true detected to all simulated rearranged markers) and precision (calculated as the ratio of true detected to all detected rearranged markers).

We observe high rates of recall and precision with small to medium block sizes and a small to medium number of events (Fig. 3A-C,E-F,I). Performance slightly decreased with increasing block sizes and number of events, probably due to the higher incidence of nested rearrangements, and a higher probability of incorrectly (but parsimoniously) assigned small blocks as rearranged. The drop in performance primarily affected syntenic moves, where it is particularly challenging to detect true rearranged blocks due to the large number of alternatives. Recall and precision remained excellent for class I nonsyntenic moves.

**Figure 3:**
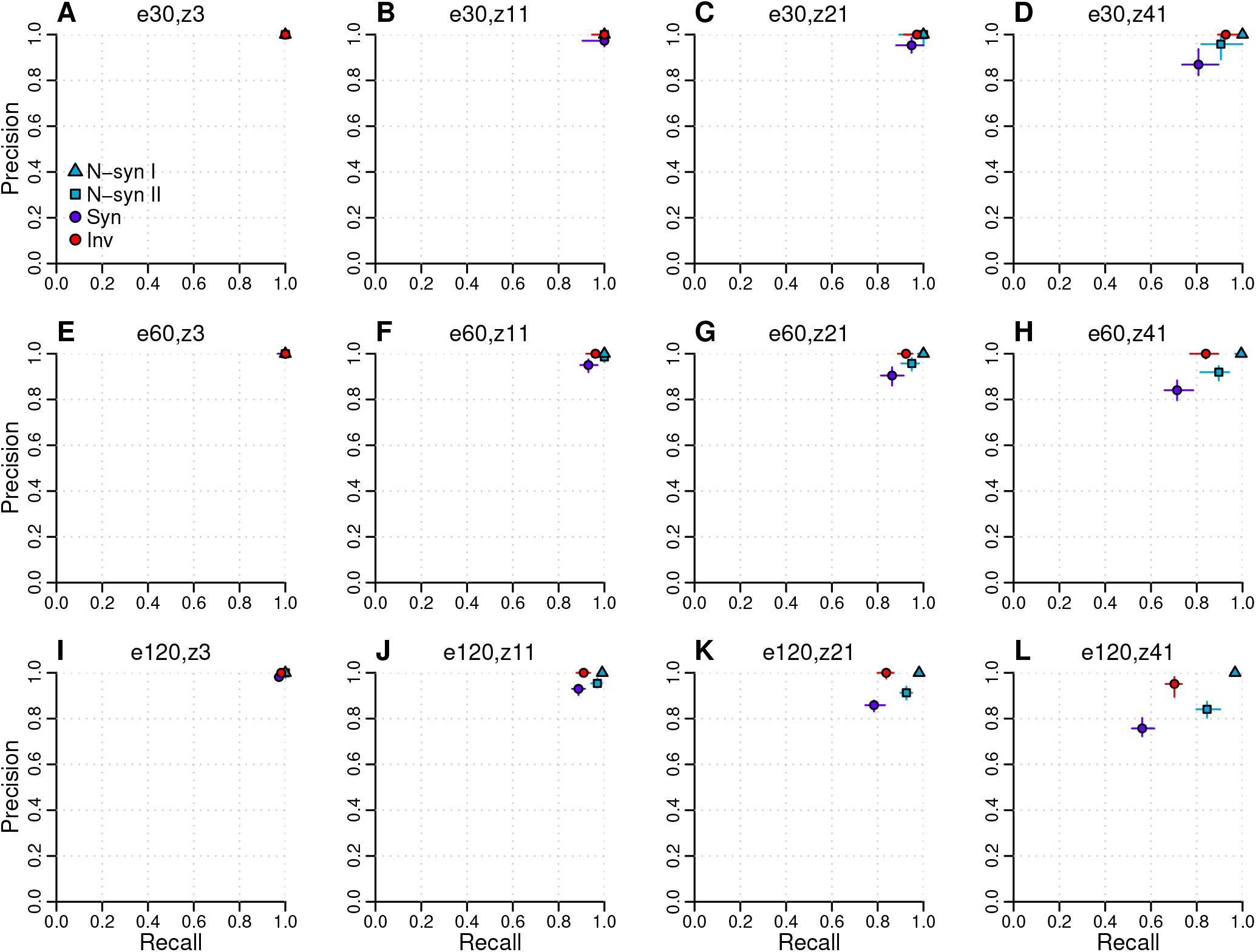
Results of performance evaluation of *rearrvisr* by simulations. Settings are shown on top of each panel (e, total number of rearrangement events; z, median block size). N-syn I, class I nonsyntenic moves; N-syn II, class II nonsyntenic moves; Syn, syntenic moves; Inv, inversions.

With a fragmented assembly of the focal genome, the performance of rearrvisr decreased only slightly (Supplementary Fig. S2A-D). With a fragmented assembly of the compared genome, recall (but not precision) decreased pronouncedly for syntenic moves, while precision (and not so much recall) decreased for class II nonsyntenic moves (Supplementary Fig. S2E-H). This shift in performance is expected, as true intrachromosomal transpositions will be more often assigned as nonsyntenic and less often as syntenic moves. Performance decreased very slightly for inversions and class I nonsyntenic moves, and only for large block sizes. With both genomes fragmented, the observed patterns combine for syntenic moves, inversions, and class II nonsyntenic moves, however we observe a distinct drop particularly in precision for class I nonsyntenic moves (Supplementary Fig. S2I-L). This effect was more pronounced with small block sizes and showed a high level of variation among runs.

We further investigated how the presence of macro-rearrangements (i.e., reciprocal translocations, fusions, and fissions) affects the detectability of rearrangements. Reciprocal translocations should be detectable as class I, but not class II nonsyntenic moves, while fusions and fissions should not be detectable by neither class I nor class II nonsyntenic moves. This is because with reciprocal translocations, FGSs are required to form circles to link matching CGSs, while fusions and fissions are not inconsistent with fragmented genome assemblies. To test this, we additionally computed recall and precision for nonsyntenic moves including or excluding markers part of macro-rearrangements (Supplementary Fig. S3). We observe that rearrvisr performs as anticipated in either case with small block sizes (Supplementary Fig. S3A,E). With increasing block size, the observed divergence between the two performance measurements diminishes, as our calculations of precision and recall are increasingly affected by markers that are simultaneously part of interchromosomal transpositions. The presence of macro-rearrangements had little effect on the identification of syntenic moves and inversions.

### 3.2 Comparison to other software

Although rearrvisr is not designed to detect the number of rearrangement events, we found that it outperforms GRIMM and UniMoG on simulated data, except for a large number of rearrangement events of very large block sizes (Fig. 4; Supplementary Figs. S4, S5). GRIMM and UniMoG, which perform similarly, consistently overestimate the number of events, particularly for fragmented genome assemblies, while rearrvisr performs well especially for a small number of events and small block sizes. Further, rearrvisr identifies balanced proportions of rearrangement classes (Fig. 4), except for fragmented compared genomes where the number of nonsyntenic moves is overestimated at the cost of syntenic moves (see above; Supplementary Fig. S4). By contrast, GRIMM tends to overestimate the relative frequency of inversions, and increasingly so with an increasing number of events and increasing block sizes.

**Figure 4:**
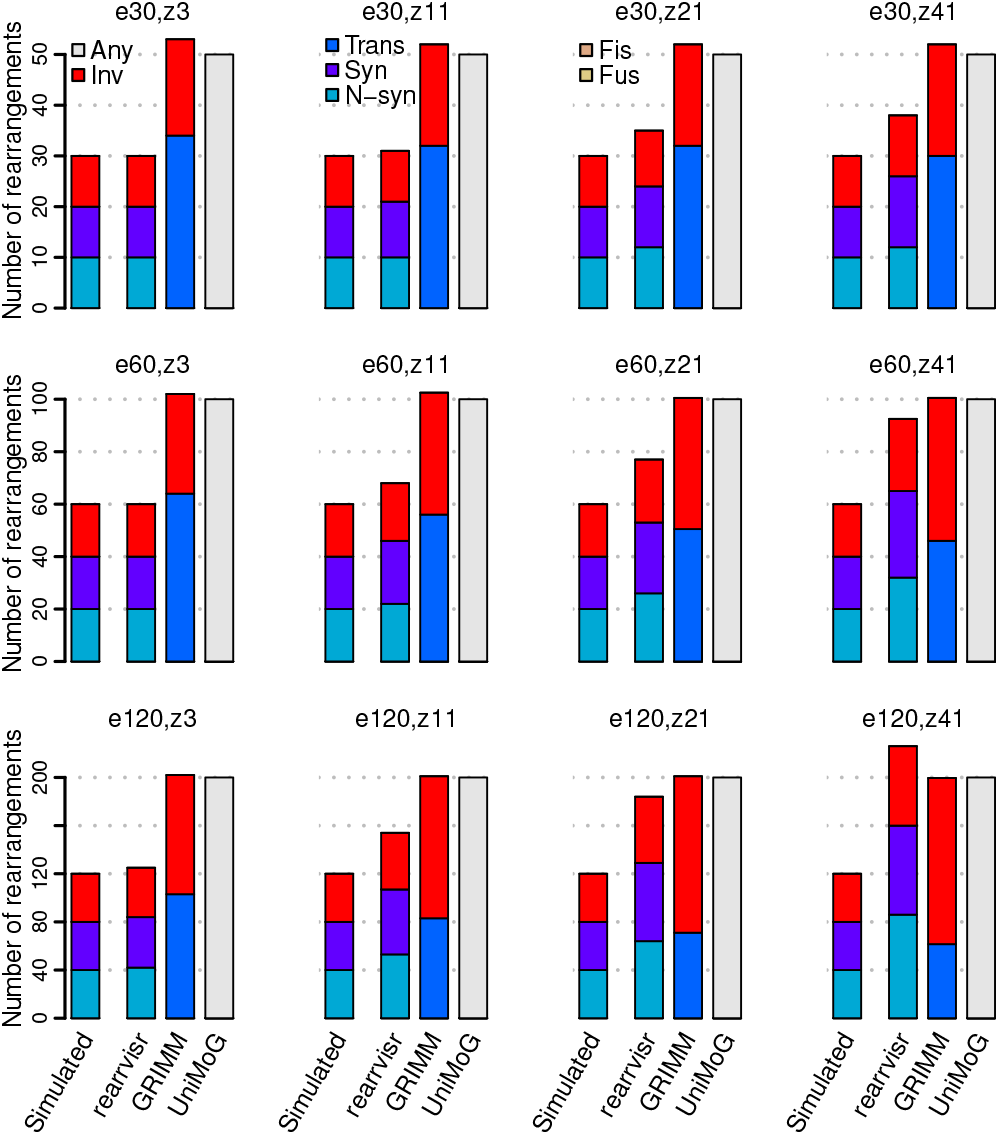
Comparison of tools by simulations. Settings are shown on top of each panel (e, total number of rearrangement events; z, median block size). Any, unclassified rearrangements; Inv, inversions; Trans, reciprocal translocations; Syn, syntenic moves; N-syn, nonsyntenic moves; Fis, fissions; Fus, fusions.

We found that GRIMM identifies the expected number of breakpoints with high accuracy (Supplementary Figs. S6, S8), except for fragmented genomes where the number of breakpoints is overestimated (Supplementary Figs. S7), as the program assumes whole biological chromosomes. By contrast, rearrvisr consistently underestimates the number of breakpoints for the majority of simulations. This should be expected, as the breakpoint of origin is not determined for nonsyntenic moves (i.e., resulting in one breakpoint per nonsyntenic move to be missing, which is roughly consistent with our results).

### 3.3 Application to Drosophila

When comparing the genomes of *D. melanogaster* and *D. simulans*, or *D. melanogaster* and *D. yakuba*, *rearrvisr* identified 16 or 8 nonsyntenic moves, 53 or 51 syntenic moves, and 19 or 51 inversions, respectively (Supplementary Table S1). GRIMM determined 26 or 15 translocations and 155 or 146 inversions, respectively. UniMoG computed 144 or 131 rearrangement events. Both *rearrvisr* and GRIMM assigned a similar number of breakpoints (200 or 204 and 212 or 211, respectively). These results are consistent with our simulations where GRIMM and UniMoG overestimate the number of rearrangement events, while *rearrvisr* slightly underestimates the number of breakpoints. When only counting rearrangements comprising three or more genes, *rearrvisr* detected 3 or 32 inversions, numbers that are in line with 1 or 27 major fixed inversions identified in previous cytological work (Lemeunier and Ashburner, 1976).

Visualizing rearrangements with the `genomeImagePlot()` function in comparison to pairwise dotplots (Supplementary Figs. S9-S12) demonstrates the ability of *rearrvisr* in projecting nonsyntenic moves and inversions onto the focal genome, and illustrates its utility in assigning syntenic moves that can be challenging to identify from dotplots. GRIMM and UniMoG project rearrangements as a sequence of transformation events onto intermediate genomes, however the generated graphics are impractical for depicting genomes with numbers of markers and rearrangements as in *Drosophila*.

In addition to comparing extant genomes, we applied *rearrvisr* to identify rearrangements between the *D. melanogaster* genome and the reconstructed ancestral genome of the *melanogaster* subgroup (Fig. 2). Our results reveal that the large inversion between *D. melanogaster* and *D. simulans* on chromosome 3R (Supplementary Figs. S9,S10) represents the derived state in *D. melanogaster*, consistent with previous work (Ranz *et al.*, 2007).

## 4 Discussion

Highly contiguous genome assemblies represent a valuable resource for detecting intra- and interspecific chromosomal rearrangements and to investigate how they contribute to adaptation, speciation, and disease. However, until now no tool was available that allowed an easy and automatable localization of rearranged markers along a genome of interest, as desirable for detecting highly rearranged regions for studies on genome structural evolution, identifying rearranged markers for correlational studies, or detecting discrepancies between different assembly versions for quality control. We thus developed *rearrvisr*, which implements an effective approach to rapidly detect and visualize rearrangements along a focal genome, facilitating the screening for structural variants. As optionally an ancestral genome reconstruction can be taken as input, rearrangements that occurred during the evolution of a specific lineage can be identified.

Our method differs fundamentally from other software designed to visualize regions of synteny by pairwise dotplots (e.g., SyMAP; Soderlund *et al.*, 2011) or dual synteny plots (e.g., GEvo; Lyons *et al.*, 2008), but which do not detect rearrangements. It also differs from tools intended to compute an optimal scenario of rearrangement events (such as GRIMM and UniMoG; reviewed in Hartmann *et al.*, 2018), but which do not assign specific markers as rearranged nor visualize rearrangements along a focal genome.

A basic assumption of our algorithm is that the provided genome maps are correct, i.e., that the assemblies or the ancestral genome reconstruction do not contain errors and that orthologous markers are accurately identified. A potential deviation from this assumption needs to be taken into account when interpreting the results. It is further required that markers are unique within each genome, i.e., genome-specific duplications are not permitted and need to be excluded from the analysis (a single paralog might be retained for tandem duplications, as then gene order is not affected). Genes or genomic regions that were duplicated before divergence of the compared genomes can be assigned as orthologs based on shared sequence similarity and can then be handled alike non-duplicated markers.

We specifically designed *rearrvisr* to work with draft genome assemblies, where unlocalized or unplaced contigs and scaffolds are permitted for the focal genome, while genome segments of the compared genome can be fragmented as long as they are contiguous. Nevertheless, we note that the performance in detecting rearrangements inherently scales with the assembly level of the input genomes. As *rearrvisr* assigns rearranged markers based on the principle of parsimony, it has limitations in correctly identifying simulated rearrangements when their number or size is large. Its primary application will thus be to genomes that differ by a modest number of small to medium-sized rearrangements.

## Acknowledgements

We thank Ivan Kryukov for valuable suggestions to improve the manuscript.

## References

Altenhoff,A.M. et al. (2015) The OMA orthology database in 2015: function predictions, better plant support, synteny view and other improvements. Nucleic Acids Research, 43, D240–D249.

Avdeyev,P. et al. (2016) Reconstruction of ancestral genomes in presence of gene gain and loss. J. Comput. Biol., 23, 150–164.

Bergeron,A. et al. (2006) A unifying view of genome rearrangements. In Bücher,P. and Moret,B.M.E. (eds) Algorithms in Bioinformatics. WABI 2006. Lecture Notes in Computer Science, Springer, Berlin, Heidelberg, Vol. 4175, pp. 163–173.

Booth,K.S. and Lueker,G.S. (1976) Testing for the consecutive ones property, interval graphs, and graph planarity using PQ-tree algorithms. J Comput. Syst. Sci., 13, 335–379.

Bhutkar,A. et al. (2008) Chromosomal rearrangement inferred from comparisons of 12 *Drosophila* genomes. Genetics, 179, 1657–1680.

Chauve,C. and Tannier,E. (2008) A methodological framework for the reconstruction of contiguous regions of ancestral genomes and its application to mammalian genomes. PLoS Comput. Biol., 4, e1000234.

Darling,A.C. et al. (2004) Mauve: multiple alignment of conserved genomic sequence with rearrangements. Genome Res., 14, 1394–1403.

Frith,M.C. and Kahn,S. (2018) A survey of localized sequence rearrangements in human DNA. Nucleic Acids Res., 46, 1661–1673.

Haas,B.J. et al. (2004) DAGchainer: a tool for mining segmental genome duplications and synteny. Bioinformatics, 20, 3643–3646.

Hannenhalli,S. and Pevzner,P.A. (1995) Transforming men into mice (polynomial algorithm for genomic distance problem). In Milwaukee,W.I. (ed.), 36th Annual Symposium on Foundations of Computer Science, IEEE Computer Soc. Press, Los Alamitos, CA, pp. 581–592.

Hartmann,T. et al. (2018) Genome rearrangement analysis: cut and join genome rearrangements and gene cluster preserving approaches. In, Setubal,J.C. et al. (eds) Comparative Genomics: Methods and Protocols. Springer New York, New York, NY, pp. 261–289.

Hilker,R. et al. (2012) UniMoG–a unifying framework for genomic distance calculation and sorting based on DCJ. Bioinformatics, 28, 2509–2511.

Hu,F. et al. (2014) MLGO: phylogeny reconstruction and ancestral inference from gene-order data. BMC Bioinformatics, 15, 354.

Jones,B.R. et al. (2012) ANGES: reconstructing ANcestral GEnomeS maps. Bioinformatics, 28, 2388–2390.

Lemeunier,F. and Ashburner,M.A. (1976) Relationships within the *melanogaster* species subgroup of the genus *Drosophila* (*Sophophora*). II. Phylogenetic relationships between six species based upon polytene chromosome banding sequences. Proc. R. Soc. B. 193, 275–294.

Li,Z. et al. (2006) Algorithmic approaches for genome rearrangement: A review. IEEE Transactions on Systems, Man, and Cybernetics, Part C (Applications and Reviews), 36, 636–648.

Lisch,D. (2013) How important are transposons for plant evolution? Nat. Rev. Gen., 14, 49–61.

Lyons,E. et al. (2008) Finding and comparing syntenic regions among *Arabidopsis* and the outgroups papaya, poplar, and grape: CoGe with rosids. Plant Physiol., 148, 1772–1781.

Ma,J. et al. (2006) Reconstructing contiguous regions of an ancestral genome. Genome Res., 16, 1557–1565.

R Core Team (2018) R: A language and environment for statistical computing. R Foundation for Statistical Computing, Vienna, Austria.

Ranz,J.M. et al. (2007) Principles of genome evolution in the *Drosophila melanogaster* species group. PLoS Biol., 5, e152.

Rieseberg,L.H. (2001) Chromosomal rearrangements and speciation. Trends Ecol. Evol., 16, 351–358.

Soderlund,C. et al. (2011) SyMAP v3.4: a turnkey synteny system with application to plant genomes. Nucleic Acids Res., 39, e68.

Tattini,L. et al. (2015) Detection of genomic structural variants from next-generation sequencing data. Front. Bioeng. Biotechnol., 3, 92.

Tesler,G. (2002) GRIMM: genome rearrangements web server. Bioinformatics, 18, 492–493.

The 1000 Genomes Project Consortium (2015) A global reference for human genetic variation. Nature, 526, 68–74.

Weischenfeldt,J. et al. (2013) Phenotypic impact of genomic structural variation: insights from and for human disease. Nat. Rev. Gen., 14, 125–138.

Yancopoulos,S. et al. (2005) Efficient sorting of genomic permutations by translocation, inversion and block interchange. Bioinformatics 21, 3340–3346.

