## Supplementary information for "*rearrvisr*: an R package to detect, classify, and visualize genome rearrangements"

### S1 Methods

#### S1.1 Simulations

To assess the performance of rearrvisr and compare it to other software, we simulated inter- and intrachromosomal transpositions (i.e., corresponding to nonsyntenic and syntenic moves, respectively), and inversions, across five chromosomes $g(i)$, $1 \leq i \leq 5$, each of size $n_i = 1000$ markers at the start of the simulation. A fixed number of rearrangement events (inter- or intrachromosomal transpositions, or inversions) occurred in random order. For each event, a source chromosome $g(i)$ was selected with probability proportional to chromosome size. A block $S$ of markers was excised from $g(i)$ between breakpoints $t_1$ and $t_2$. The first breakpoint $t_1$ was drawn from a uniform distribution $\alpha_i \sim \mathrm{Uniform}(1, n_i - 1)$, and the second breakpoint $t_2$ was assigned at $z + 1$ markers distance up- or downstream from $t_1$ (potentially truncating $S$ at the chromosome end). The value informing block size, $z$, was drawn from a negative binomial distribution with parameters size $\eta$ and probability $\varphi$. For an interchromosomal transposition, a destination chromosome $g(j) \neq g(i)$ was selected with probability proportional to chromosome size and a third breakpoint $t_2$ on $g(j)$ was drawn from a uniform distribution $\beta_j \sim \mathrm{Uniform}(0, n_j + 1)$. Block $S$ was then inserted at $t_3$ on $g(j)$. For an intrachromosomal transposition, $t_3$ on $g(i)$ was drawn from a uniform distribution $\beta_i \sim \mathrm{Uniform}(0, n_i + 1)$ and requiring $t_3$ being at a distance of at least one marker up- or downstream of $S$. Block $S$ was then inserted at $t_3$. For an inversion, block $S$ was inserted in reversed orientation on $g(i)$. We simulated small ($\eta = 20$; $\varphi = 0.92$; median $z = 2$), medium ($\eta = 15$; $\varphi = 0.6$; median $z = 10$), large ($\eta = 10$; $\varphi = 0.325$; median $z = 20$), or very large ($\eta = 6$; $\varphi = 0.123$; median $z = 40$) block sizes, equal numbers of rearrangement classes with a total of 30, 60, or 120 rearrangement events, and ran 100 replicates per setting.

In addition, we tested how genome fragmentation affects the detectability of rearrangements. For this, a randomly selected genome segment $g(i) \in G$ was split at marker $g(i)_m$, $1 \leq m \leq n_i$, into $g(i_1) = (g(i)_1, \ldots, g(i)_m)$ and $g(i_2) = (g(i)_{m+1}, \ldots, g(i)_{n_i})$. The index $m$ was slightly skewed to either end of $g(i)$ by assigning $m = \mathrm{floor}(t_i(n_i - 1) + 1)$, where $t_i$ was drawn from a beta distribution with shape parameters 1.5 and 8. Segment $g(i)$ was replaced by $g(i_1)$ and $g(i_2)$ on $G$, forming the new set of segments $g(i) \in G$, and the process repeated to introduce a total of $s = 20$ splits.

Scripts used to perform the simulations are available on GitHub at https://github.com/dorolin/rearrvisr_simulations.

#### S1.2 *Drosophila* data

##### S1.2.1 Data preparation

Peptide sequences and annotation information (`pep.all.fa` and `gff3` files) for genes from 12 *Drosophila* species were downloaded on Dec 23 2017 from Ensembl Release 91 (http://dec2017.

archive.ensembl.org; *D. melanogaster*) or Ensembl Metazoa Release 37 (http://oct2017-metazoa.ensembl.org; *D. ananassae*, *D. erecta*, *D. grimshawi*, *D. mojavensis*, *D. persimilis*, *D. pseudoobscura*, *D. sechellia*, *D. simulans*, *D. virilis*, *D. willistoni*, and *D. yakuba*). Sequences that were shorter than 50 amino acids or contained premature stop codons were excluded. 20,803 orthologous groups were identified with OMA standalone v2.2.0 (Altenhoff *et al.*, 2015), using as guidance tree the phylogeny published in Drosophila 12 Genomes Consortium (2007), and default settings otherwise. A single representative splicing variant per gene was identified by OMA, and alternative splicing variants were excluded from subsequent analyses. The retained number of genes per species were 15,052, 14,998, 14,969, 13,818, 14,581, 16,856, 15,845, 16,409, 15,355, 14,477, 15,490, and 16,039 for *D. ananassae*, *D. erecta*, *D. grimshawi*, *D. melanogaster*, *D. mojavensis*, *D. persimilis*, *D. pseudoobscura*, *D. sechellia*, *D. simulans*, *D. virilis*, *D. willistoni*, and *D. yakuba*, respectively.

Sequences of 4,792 OMA orthologous groups that only included one-to-one orthologous genes present in all 12 species were extracted and individually aligned with MAFFT v7.407 (Katoh *et al.*, 2002; Katoh and Standley, 2013), using the iterative refinement method incorporating local pairwise alignment information (`--localpair --maxiterate 1000` settings). Alignments with not more than 20% missing data (4,308 orthologous groups) were concatenated, and a phylogenetic tree was computed with RAxML v8.2.12 (Stamatakis, 2014). The best protein substitution model (JTT with empirical base frequencies) was determined by the program (`-f a -# autoMRE -m PROTGAMMAAUTO --auto-prot=ml` settings). The resulting tree was rooted at the branch that best balanced the subtree lengths.

##### S1.2.2 Ancestral genome reconstruction

Genome maps were prepared for all retained genes (i.e., excluding low quality sequences and non-representative splicing variants) based on gene position information extracted from the `gff3` files. Gene start and end positions were calculated as the average of CDS midpoints -/+ 1 base pair to avoid the occurrence of overlapping gene positions, which are not supported by the genome reconstruction software ANGES v1.01 (Jones *et al.*, 2012). A few remaining overlaps between gene positions were resolved manually. The ANGES `marker` input file was generated using the genome maps of all 12 *Drosophila* species and 20,803 OMA orthologous groups (from the OMA `OrthologousGroups.txt` file) with the R script `oma2anges.R` (available in the `exec` directory of the *rearrvisr* package). The ANGES `tree` input file was based on the best rooted RAxML tree computed above (including branch lengths). The ancestral genome of the *melanogaster* subgroup *'MSSYE'* (Drosophila 12 Genomes Consortium, 2007; i.e., separating *D. melanogaster*, *D. simulans*, *D. sechellia*, *D. yakuba*, and *D. erecta* from the remainder of the *Drosophila* species) was reconstructed using the ANGES master pipeline (`anges_CAR.py`) and options `markers_doubled 1` (infer ancestral marker orientation), `markers_unique 2` (no duplicated markers), `markers_universal 1` (no missing markers in ingroup), `c1p_telomeres 0` (no telomeres), and `c1p_heuristic 1` (using a greedy heuristic).

A total of 27,242 *Ancestral Contiguous Sets* (ACS; Jones *et al.*, 2012) were identified by ANGES, of which 26,594 ACS were organized into 20 CARs (648, or 2.4%, of ACS were discarded by the program). These CARs comprised a total of 8,973 ancestral markers, and 99.2% of them were grouped into four major CARs (i.e., the 20 CARs included 4,250, 1,914, 1,711, 1,026, 28, 23, 5, 3, 2, and 11 x 1 ancestral markers).

Scripts used to generate the *Drosophila* data set are available on GitHub at https://github.com/dorolin/rearrvisr_dataprep.

### Tables

Table S1: Numbers of rearrangements and breakpoints detected between *D. melanogaster* and *D. simulans* (*mel-sim*) or *D. melanogaster* and *D. yakuba* (*mel-yak*).

| | **rearrvisr** | | **GRIMM** | | **UniMoG** | |
|---|---|---|---|---|---|---|
| | *mel-sim* | *mel-yak* | *mel-sim* | *mel-yak* | *mel-sim* | *mel-yak* |
| Nonsyntenic moves | 16 | 8 | – | – | – | – |
| Syntenic moves | 53 | 51 | – | – | – | – |
| Translocations | – | – | 26 | 15 | – | – |
| Inversions | 19 | 52 | 155 | 146 | – | – |
| **Total rearrangements** | 88 | 111 | 181 | 161 | 144 | 131 |
| Internal breakpoints | – | – | 210 | 209 | – | – |
| External breakpoints | – | – | 2 | 2 | – | – |
| **Total breakpoints** | 200 | 204 | 212 | 211 | – | – |

#### Figures

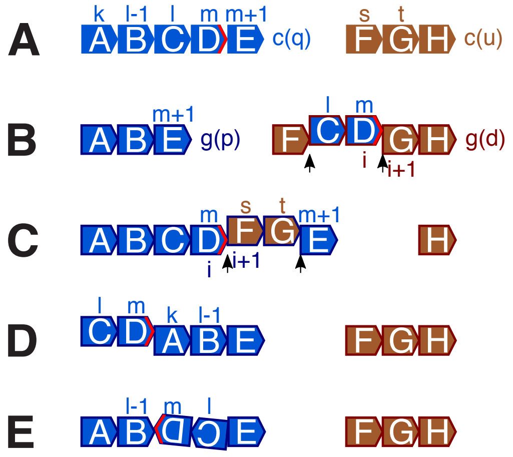


**Figure S1:** Oriented markers are illustrated with pentagons, ordered on their GSs. **(A)** Markers A–E are on CGS $c(q)$ and markers F-H are on CGS $c(u)$. Indices above markers denote their positions on their CGSs (see main text). The tail of marker $c(q)_m$ (marker D) is indicated in red. **(B–E)** Markers that are orthologs to A–H on $c(q)$ and $c(u)$ are rearranged on GS $g(p)$ or GS $g(d)$. Rearrangements are defined through the formation of a novel adjacency of marker $c(q)_m$, here illustrated with the tail of marker D. Novel internal boundaries formed through nonsyntenic moves are indicated with black arrowheads. Indices below markers denote their positions on $g(p)$ and $g(d)$ after the occurrence of rearrangements. **(B)** A nonsyntenic move resulting from the movement of a block of markers from $c(q)$ onto $g(d)$, resulting in the formation of novel internal boundaries of $c(q)$ on $g(d)$. **(C)** A nonsyntenic move resulting from the movement of a block of markers from $c(u)$ onto $g(p)$, resulting in the formation of novel internal boundaries of $c(q)$ on $g(p)$. **(D)** A syntenic move. **(E)** An inversion.

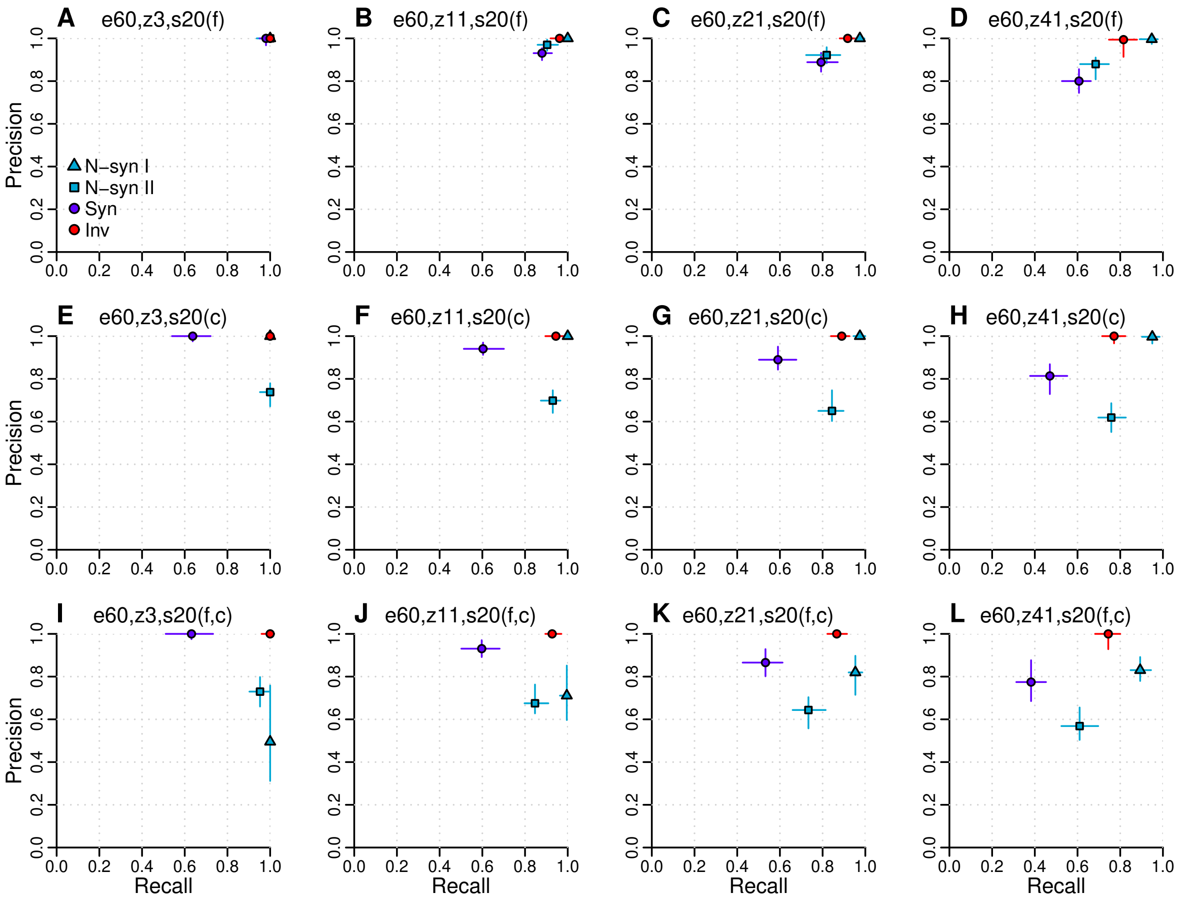


**Figure S2:** Results of performance evaluation of *rearrvisr* by simulations for runs with genome fragmentation. Settings are shown on top of each panel (e, total number of rearrangement events; z, median block size; s, number of introduced splits for the genomes in parentheses: f, focal genome; c, compared genome). N-syn I, class I nonsyntenic moves; N-syn II, class II nonsyntenic moves; Syn, syntenic moves; Inv, inversions.

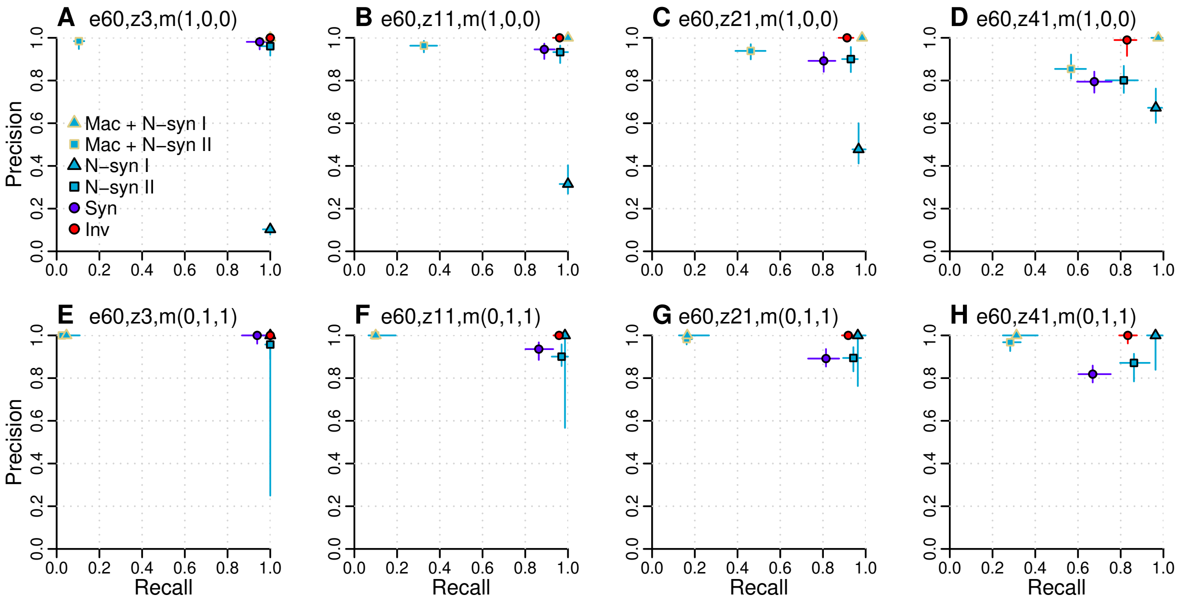


**Figure S3:** Results of performance evaluation of *rearrvisr* by simulations for runs with macro-rearrangements. Settings are shown on top of each panel (e, total number of rearrangement events; z, median block size; m, number of additional macro-rearrangements in parentheses: reciprocal translocation,fission,fusion). N-syn I, class I nonsyntenic moves; N-syn II, class II nonsyntenic moves; Syn, syntenic moves; Inv, inversions.

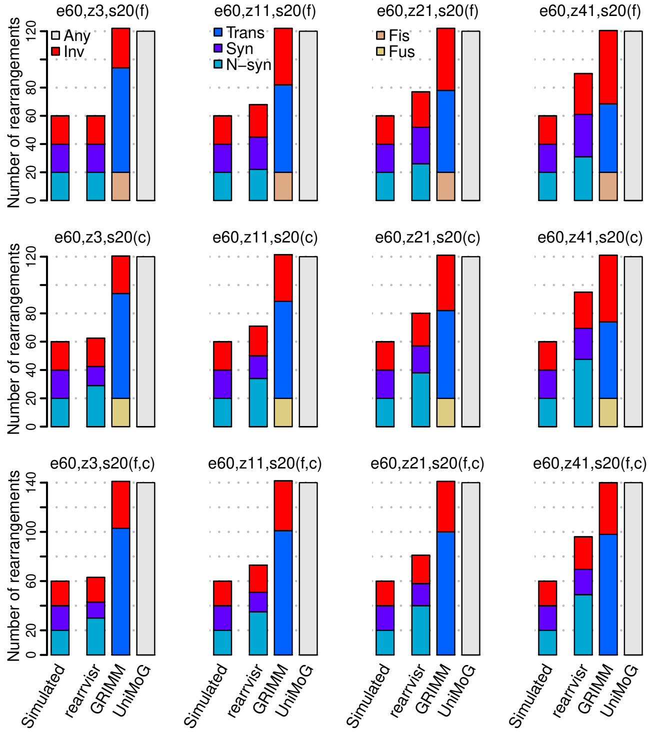


**Figure S4:** Comparison of tools by simulations for runs with genome fragmentation. Settings are shown on top of each panel (e, total number of rearrangement events; z, median block size; s, number of introduced splits for the genomes in parentheses: f, focal genome; c, compared genome). Any, unclassified rearrangements; Inv, inversions; Trans, reciprocal translocations; Syn, syntenic moves; N-syn, nonsyntenic moves; Fis, fissions; Fus, fusions.

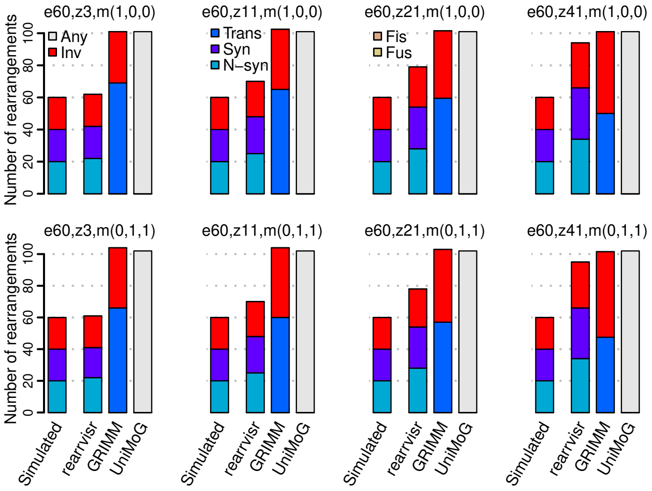


**Figure S5:** Comparison of tools by simulations for runs with macro-rearrangements. Settings are shown on top of each panel (e, total number of rearrangement events; z, median block size; m, number of additional macro-rearrangements in parentheses: reciprocal translocation,fission,fusion). Any, unclassified rearrangements; Inv, inversions; Trans, reciprocal translocations; Syn, syntenic moves; N-syn, nonsyntenic moves; Fis, fissions; Fus, fusions.

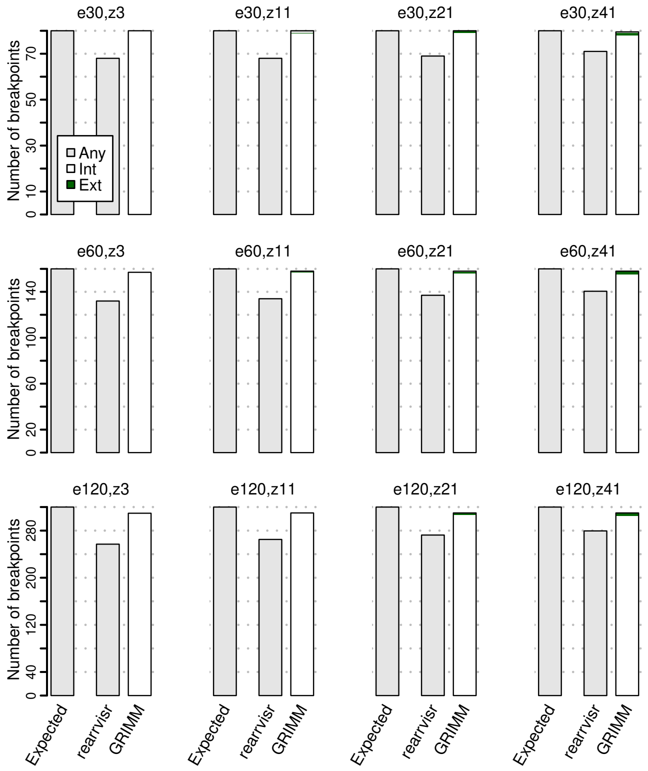


**Figure S6:** Comparison of tools by simulations. Simulation settings are shown on top of each panel (e, total number of rearrangement events; z, median rearrangement size). Any, unclassified breakpoints; Int, internal breakpoints; Ext, external breakpoints.

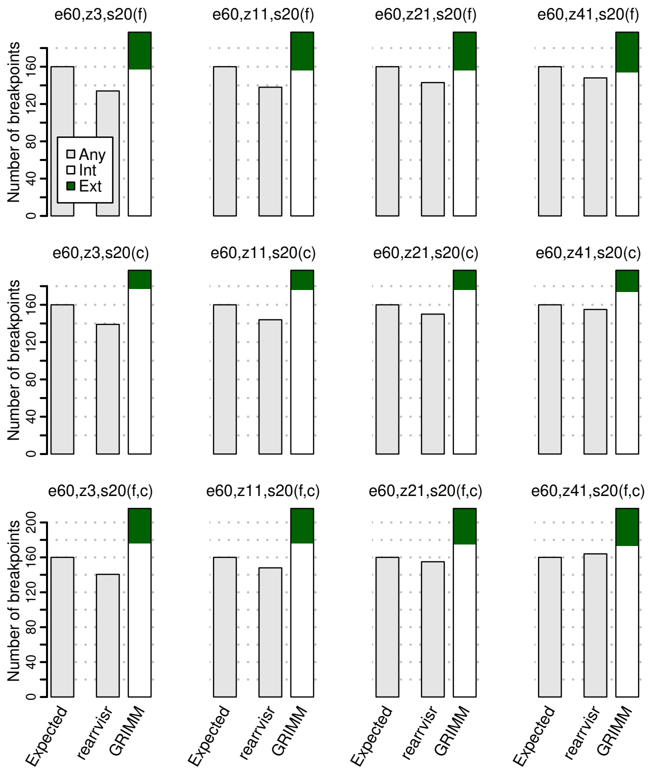


**Figure S7:** Comparison of tools by simulations for runs with genome fragmentation. Simulation settings are shown on top of each panel (e, total number of rearrangement events; z, median rearrangement size; s, number of introduced splits for the genomes in parentheses: f, focal genome; c, compared genome). Any, unclassified breakpoints; Int, internal breakpoints; Ext, external breakpoints.

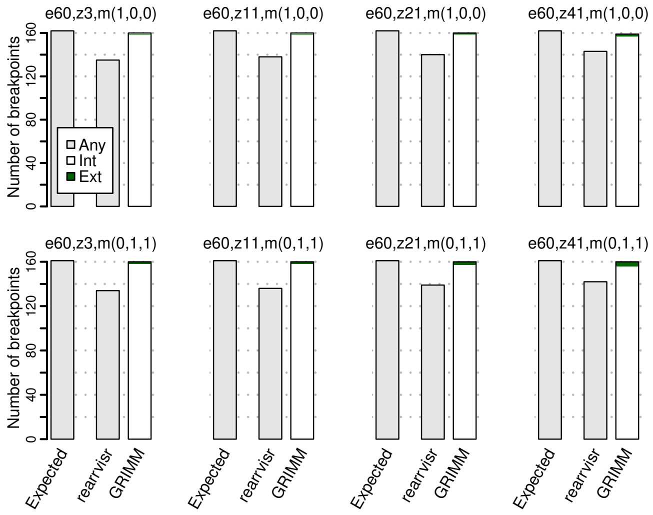


**Figure S8:** Comparison of tools by simulations for runs with macro-rearrangements. Simulation settings are shown on top of each panel (e, total number of rearrangement events; z, median rearrangement size; m, number of additional macro-rearrangements in parentheses: reciprocal translocation,fission,fusion). Any, unclassified breakpoints; Int, internal breakpoints; Ext, external breakpoints.

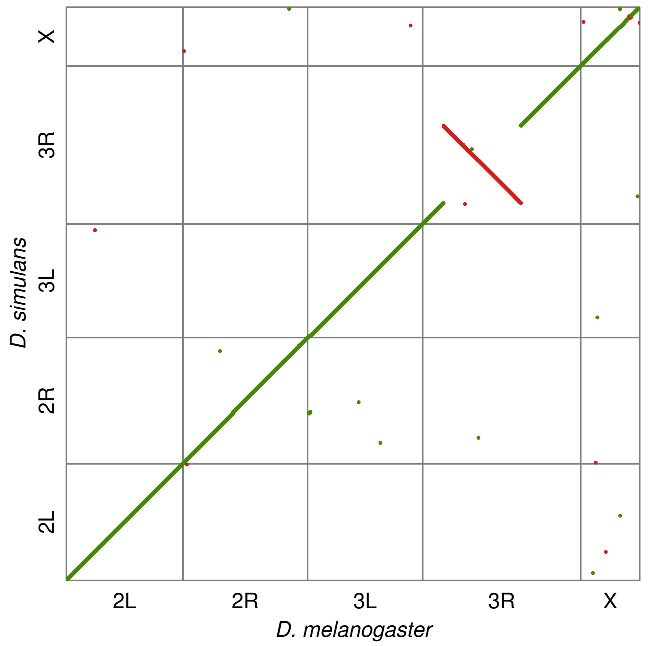


**Figure S9:** Dotplot *D. melanogaster* vs. *D. simulans*.

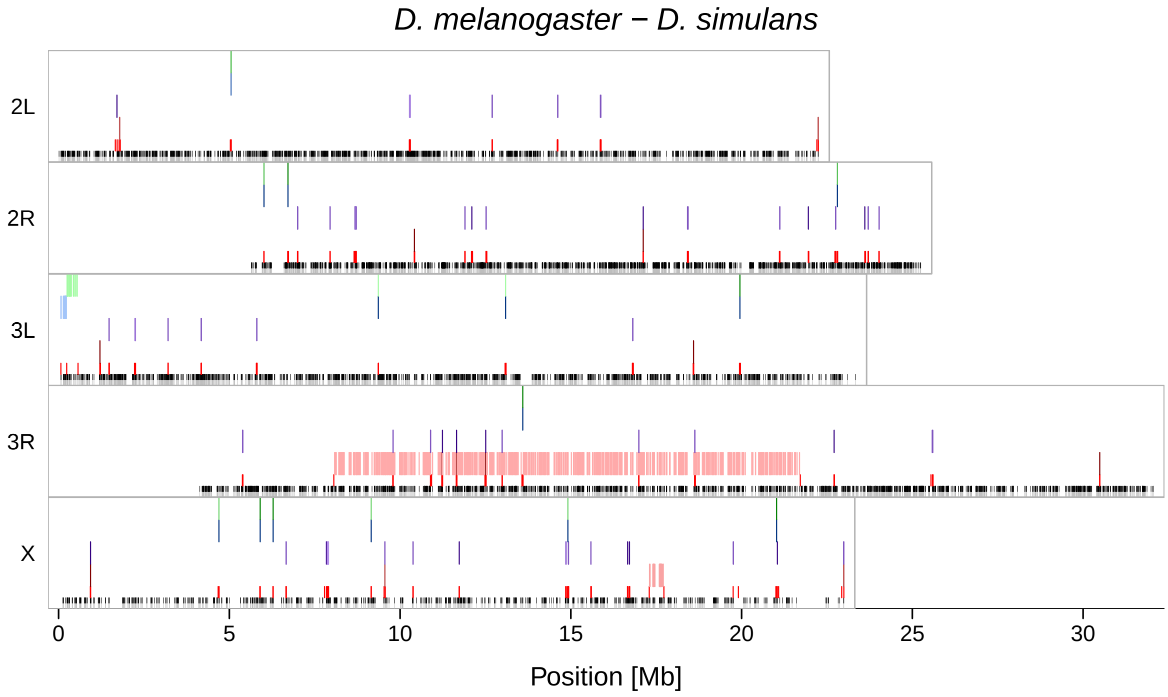


**Figure S10:** Output of the `genomeImagePlot()` function with *D. melanogaster* as focal genome and *D. simulans* as compared genome.

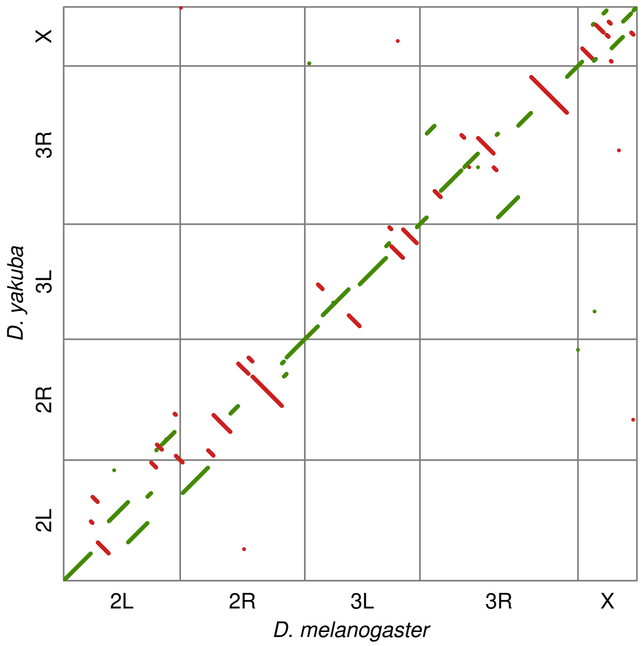


**Figure S11:** Dotplot *D. melanogaster* vs. *D. yakuba*.

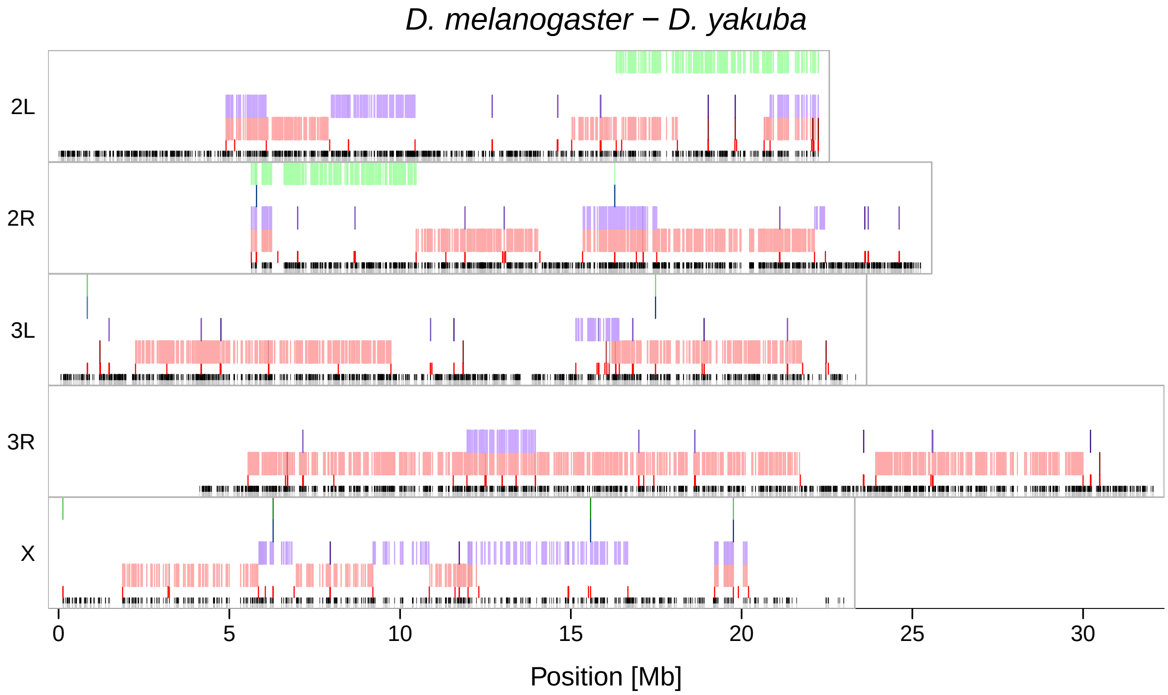


**Figure S12:** Output of the `genomeImagePlot()` function with *D. melanogaster* as focal genome and *D. yakuba* as compared genome.

### References

Altenhoff,A.M. *et al*. (2015) The OMA orthology database in 2015: function predictions, better plant support, synteny view and other improvements. *Nucleic Acids Research*, **43**, D240–D249.

Drosophila 12 Genomes Consortium (2007) Evolution of genes and genomes on the *Drosophila* phylogeny. *Nature*, **450**, 203-218.

Jones,B.R. *et al*. (2012) ANGES: reconstructing ANcestral GEnomeS maps. *Bioinformatics*, **28**, 2388-2390.

Katoh,K. and Standley,D.M. (2013) MAFFT multiple sequence alignment software version 7: improvements in performance and usability. *Molecular Biology and Evolution*, **30**, 772–780.

Katoh,K. *et al*. (2002) MAFFT: a novel method for rapid multiple sequence alignment based on fast Fourier transform. *Nucleic Acids Research*, **30**, 3059–3066.

Stamatakis,A. (2014) RAxML version 8: a tool for phylogenetic analysis and post-analysis of large phylogenies. *Bioinformatics*, **30**, 1312–1313.
